# Harmful Algal Bloom-Forming Organism Responds to Nutrient Stress Distinctly From Model Phytoplankton

**DOI:** 10.1101/2021.02.08.430350

**Authors:** Craig McLean, Sheean T. Haley, Gretchen J. Swarr, Melissa C. Kido Soule, Sonya T. Dyhrman, Elizabeth B. Kujawinski

## Abstract

- Resources such as nitrogen (N) and phosphorus (P) play an important role in primary production and constraining phytoplankton bloom dynamics. Models to predict bloom dynamics require mechanistic knowledge of algal metabolic shifts in response to resource limitation. For well-studied model phytoplankton like diatoms, this information is plentiful. However, for less-studied groups such as the raphidophytes, there remain significant gaps in understanding metabolic changes associated with nutrient limitation.
- Using a novel combination of metabolomics and transcriptomics, we examined how the harmful algal bloom-forming raphidophyte *Heterosigma akashiwo* shifts its metabolism under N- and P-stress. We chose *H. akashiwo* because of its ubiquity within estuarine environments worldwide, where bloom dynamics are influenced by N and P availability.
- Our results show that each stress phenotype is distinct in both the allocation of carbon and the recycling of macromolecules. Further, we identified biomarkers of N- and P-stress that may be applied *in situ* to help modelers and stakeholders manage, predict, and prevent future blooms.
- These findings provide a mechanistic foundation to model the metabolic traits and trade-offs associated with N- and P-stress in *H. akashiwo*, and evaluate the extent to which these metabolic responses can be inferred in other phytoplankton groups.

## Introduction

The scarcity of nitrogen (N) and phosphorus (P) limits primary production across aquatic ecosystems (Moore *et al.*, 2013; Ardyna & Arrigo, 2020) by regulating the growth and structure of phytoplankton communities (Arrigo, 2005; Follows *et al.*, 2007; Harke *et al.*, 2016). These communities consist of an extremely diverse group of phylogenetically and physiologically distinct phytoplankton (Keeling *et al.*, 2014; Caron *et al.*, 2017). The abundance of individual phytoplankton groups can vary widely over space and time (de Vargas *et al.*, 2015), due in large part to their group-specific evolutionary and ecological strategies for managing resource limitation. These strategies vary from reallocation of intracellular nutrients (Van Mooy *et al.*, 2009; Allen *et al.*, 2011; Kujawinski *et al.*, 2017), reduction of nutrient quotas needed for growth (Grzymski & Dussaq, 2012; Read *et al.*, 2017), or increased extracellular scavenging (Haley *et al.*, 2017; Harke *et al.*, 2017; Kujawinski *et al.*, 2017).

The impact of distinct nutrient response strategies is most evident when considering harmful algal blooms (HABs). HABs may occur when a set of physiological traits allow a single phytoplankton group to out-compete its neighbors (Wilson *et al.*, 2019). Such blooms have increased in frequency over the past 40 years with climate change and eutrophication (Wells *et al.*, 2015), causing hundreds of millions of dollars in economic damages to fisheries and public health (Hoagland *et al.*, 2002). Many blooms have been linked to resource availability (Anderson *et al.*, 2008), and as a result, knowledge of nutrient response mechanisms and their associated trade-offs towards fitness holds tremendous promise in managing HABs (Sharma & Steuer, 2019). The mechanics of how nutrients influence metabolic response strategies have been identified in only a few model phytoplankton (*e.g.*, diatoms (Brembu *et al.*, 2017), coccolithophores (Rokitta *et al.*, 2014; Rokitta *et al.*, 2016; Wördenweber *et al.*, 2018)), leaving significant gaps in our understanding of nutrient responses in HAB-forming groups.

We do not know the extent to which metabolism data from model phytoplankton reflects that of non-model phytoplankton. Although core metabolic functional redundancies occur in all primary producers (Louca *et al.*, 2018), physiological studies suggest that phytoplankton groups vary widely in their capabilities beyond carbon processing (Smayda, 1997). For example, elemental stoichiometry studies observe that phytoplankton groups differ in their intracellular macromolecular pool composition (Bonachela *et al.*, 2016), and transcriptome studies show that phytoplankton respond differently to environmental perturbations (Alexander *et al.*, 2015b). Therefore, direct evaluations of non-model phytoplankton metabolism, specifically under conditions of acute shortages of essential nutrients (stress), are needed to evaluate the extent to which metabolic knowledge from model phytoplankton can describe other groups.

One factor driving the knowledge discrepancy between model and non-model phytoplankton groups is the paucity of fully sequenced eukaryotic genomes for most phytoplankton, which are required to build genome-scale models of metabolism (Thiele & Palsson, 2010; Levitan *et al.*, 2014; Smith *et al.*, 2019). Due to this constraint, investigators have used ‘omics techniques like transcriptomics or metabolomics to characterize metabolic responses of phytoplankton to nutrient stress (Wurch *et al.*, 2011; Rokitta *et al.*, 2014; Alipanah *et al.*, 2015; Rokitta *et al.*, 2016; Brembu *et al.*, 2017; Alipanah *et al.*, 2018; Hennon & Dyhrman, 2019; Wurch *et al.*, 2019). While each of these methods offers substantial insights on physiological differences among phytoplankton (Alexander *et al.*, 2015a), they fail to capture the system-level dynamics driving changes in metabolism when used in isolation (Wördenweber *et al.*, 2018). For example, while metabolomic approaches provide evidence of biochemical reactions, predicting the mechanism driving the activity is challenging. By contrast, transcriptomic techniques reveal pathway level changes, yet such changes may be inhibited by post-translational regulatory processes beyond the scope of the data. Applying metabolomic and transcriptomic methods in tandem circumvents these issues. However, computational challenges have typically limited these multi-‘omic efforts to targeted analyses of specific pathways rather than system-level changes (Kujawinski *et al.*, 2017). Combining ‘omics techniques has tremendous promise for understanding the diversity of phytoplankton responses to N- and P-stress.

In this study, we examined N- and P-stress metabolism using a combination of metabolomics and transcriptomics data for the HAB-forming raphidophyte, *Heterosigma akashiwo*. *H. akashiwo* populations are distributed ubiquitously within subtropical environments (Taylor & Haigh, 1993; Smayda, 1998), and their blooms have caused significant economic losses (Rensel, 2007). Both N- and P-stress are known to be important drivers of *H. akashiwo* blooms (Shikata *et al.*, 2008; Ji *et al.*, 2018). Our findings provide a mechanistic foundation to manage, predict, and prevent blooms of *H. akashiwo* and suggest that a broader understanding of non-model phytoplankton metabolism is necessary to understand how phytoplankton communities will adapt to a changing ocean.

## Method

### Culture Maintenance

We cultured *H. akashiwo* strain CCMP 2393 (isolated from Rehoboth Bay, Delaware, USA) in L1 medium (882 mM NaNO_3_, 36.2 mM NaH_2_PO_4_) made with autoclaved 0.2-μm filtered seawater from Vineyard Sound, MA. Cultures were not axenic, but were uni-algal and uni-eukaryotic. We grew each culture with light intensity of 100 μmol quanta m^−2^ s^−1^ of photosynthetically active radiation (400-700 nm) during a 14h:10h light:dark cycle at 18°C.

### Experimental Design

We used entrainment cultures to decrease carryover of nutrients from stock cultures and promote acclimation to the experimental conditions. We grew single entrainment cultures (~100 mL) in modified L1 medium (base seawater as above) under the following nitrogen and phosphorus conditions: replete (576 μM NaNO_3_ and 36.2 μM NaH_2_PO_4_), N-stress (5 μM NaNO_3_ and 36.2 μM NaH_2_PO_4_), and P-stress (5 μM NaNO_3_ and 0.2 μM NaH_2_PO_4_). We kept entrainment cultures at 18 °C on a 14:10 light:dark cycle (as above) with gentle rotation (75 rpm) for 3 days. We initiated experiments by inoculating triplicate 1-L flasks (containing 0.3 L media) for each treatment with 30 mL of the corresponding entrainment culture. We maintained experimental flasks under the same conditions as the entrainment cultures.

We monitored growth in each flask by *in vivo* chlorophyll fluorescence on a Turner Designs Aquafluor handheld fluorometer with paired cell counts. We preserved cell count samples in 2% (final concentration) acid Lugol’s solution and we determined cell concentrations by microscopy. We took cell concentration measurements at the same time each day (during the middle of the light phase) to avoid diel changes in metabolite synthesis. We harvested replete cultures in exponential phase, and harvested N-stress and P-stress cultures once growth rates and cell yields were reduced relative to the replete control, in agreement with the definition of nutrient stress rather than deficiency (MacIntyre & Cullen, 2005). Specifically, we harvested treatments for metabolomics analysis on day 3, when we observed significant differences (p < 0.001, Tukey-HSD Test, n = 3) in cell counts between stressed and replete cultures (Fig S1). We filtered cells (300 mL) of each replicate in each treatment onto combusted 47 mm GF/Fs (Whatman) using combusted glass filtration funnels under low vacuum pressure (never exceeding 5 mm Hg) to collect particulate metabolite samples. We flash-froze filters in cryovials in liquid nitrogen and stored them at −80°C until extraction.

### Filter Extractions

We split each filter in half for separate extractions for targeted and untargeted metabolomic analyses. The extraction procedure for untargeted and targeted samples are identical with the exception of the final solid phase extraction (see below). For both types of analyses, we first cut each filter half into six roughly equivalent pieces and placed them into an 8-mL amber glass vial. We extracted metabolites from filters using cold 40:40:20 acetonitrile: methanol:water+0.1 M formic acid similar to previous work (Rabinowitz & Kimball, 2007; Kido Soule *et al.*, 2015). We then added 25 μL of 1 μg/mL deuterated standard mix (d_3_-glutamic acid, d_4_-4-hydroxybenzoic acid, and d_5_-taurocholate) as extraction recovery standards. We sonicated the solvent-filter mixture for 10 minutes to lyse the cells, and transferred the solvent into a microcentrifuge tube. We rinsed the filters with three 200-μL aliquots of extraction solvent to capture any remaining organic matter. We centrifuged the combined extracts at 20,000 × *g* for 5 minutes, and transferred the supernatant into clean 8-mL amber glass vials with care to leave behind any filter or cellular debris. We neutralized the extracts with 25.6 μL of 6 M ammonium hydroxide and dried them down to near dryness in a vacufuge. We reconstituted dried samples for targeted analysis in 95:5 water:acetonitrile solution plus 2.5 μL of 5 μg/mL deuterated standard mix (d3-glutamic acid, d4-4-hydroxybenzoic acid, d5-taurocholate).

For the untargeted analysis, a PPL extraction step is necessary. We reconstituted these samples with 500 μL 0.05 μg/mL d_2_-biotin with 0.01 M HCl to lower the pH to 2 and ran these samples through 100 mg/1 mL Agilent Bond Elut PPL cartridges. We pre-conditioned the cartridge with one cartridge-volume of 100% methanol and passed acidified untargeted samples through the cartridge at a flow rate below 40 mL min^−1^. We rinsed the cartridges with one cartridge-volume of 0.01 M HCl, dried them down for 5 minutes, and eluted the metabolites with 1 mL of methanol. We dried untargeted samples again to near dryness and reconstituted them with 250 μL of 95:5 water:acetonitrile plus 2.5 μL of 5 μg/mL deuterated standard mix (d3-glutamic acid, d4-4-hydroxybenzoic acid, d5-taurocholate). We combined 45 μL aliquots from each sample to create a pooled sample.

### Liquid Chromatography and Mass Spectrometry

We analyzed metabolite samples for untargeted analyses by high-performance liquid chromatography (HPLC, Micro AS autosampler and Surveyor MS Pump Plus, Thermo Scientific) coupled via electrospray ionization (ESI) to a hybrid linear ion trap-Fourier transform ion cyclotron resonance (FT-ICR) mass spectrometer (7T LTQ FT Ultra, Thermo Scientific). We separated metabolites on a Synergi Fusion reverse phase C_18_ column (4 μm, 2.0 x 150 mm, Phenomenex), equipped with a guard column and precolumn filter, and maintained at 35°C. We eluted the column with (A) 0.1% formic acid in water and (B) 0.1% formic acid in acetonitrile at a flow rate of 0.25 mL min^−1^. We held the column at 5% B for 2 min, ramped to 65% B over 18 min, quickly ramped to 100% B over 5 min, held at 100% B for 7 min and then equilibrated at 5% B for 8 min prior to the next injection (total run time = 40 min). We injected 20 μL of sample individually for positive and negative ion mode analyses. We externally calibrated the mass spectrometer just prior to analysis in positive and negative ion modes using the manufacturer’s solutions. We optimized the capillary temperature and ESI voltage at 330°C and 4.2 kV in positive mode and at 365°C and 3.8 kV in negative mode. We maintained sheath gas, auxiliary gas, and sweep gas flow rates at 35, 5, and 2, respectively (arbitrary units) for both polarities. We collected MS and data dependent MS/MS scans as follows: (1) a full MS scan in the FT-ICR analyzer from 100-1000 *m/z*, with mass resolving power set to 100,000 (defined at *m/z* 400); and (2) collision-induced dissociation fragmentation scans (MS/MS) in the linear ion trap for the four most abundant ions in each full scan. We collected MS/MS spectra under dynamic exclusion with an exclusion time of 20 seconds. At the start of each batch, we injected the pooled sample multiple times to condition the column with the sample matrix and to stabilize peak retention times. We also analyzed the pooled sample every nine samples for quality assurance.

We analyzed targeted samples by ultrahigh-performance liquid chromatography (UHPLC, Accela Open Autosampler and Accela 1250 Pump, Thermo Scientific) coupled via heated electrospray ionization (H-ESI) to a triple quadrupole mass spectrometer (TSQ Vantage, Thermo Scientific) operated under selected reaction monitoring (SRM) mode. We set the spray voltage at 4000 V (positive mode) and 3200 V (negative mode). We set source gases at 55 (sheath) and 20 (aux gas), heated capillary temperature at 375 °C, and the vaporizer temperature at 400 °C. We performed chromatographic separation on a Waters Acquity HSS T3 column (2.1 × 100 mm, 1.8 μm) equipped with a Vanguard pre-column and maintained at 40 °C. We eluted the column with (A) 0.1% formic acid in water and (B) 0.1% formic acid in acetonitrile at a flow rate of 0.5 mL min^−1^. The gradient starts at 1% B for 1 min, ramp to 15% B from 1-3 min, ramp to 50% from 3-6 min, ramp to 95% B from 6-9 min, hold until 10 min, ramp to 1% B from 10-10.2 min, with final re-equilibration at 1% B (total gradient time 12 min). We made separate autosampler injections of 5 μL for positive and negative ion modes.

### Standard Optimization

We obtained authentic standards at the highest grade available from Sigma Aldrich for compounds outside of our existing targeted method (Johnson *et al.*, 2017). We injected standards at concentrations of 1 μg/mL in Milli-Q water to optimize selected reaction monitoring (SRM) conditions (s-lens, collision energy, product ions). We monitored at least two SRM transitions (precursor-product ion pairs) for quantification and confirmation of each target compound, derived from optimization protocols that maximize analyte signal-to-noise. We determined the chromatographic retention time of each compound with standards dissolved in Milli-Q.

### Data Processing

We converted untargeted data files from proprietary Thermo RAW into mzML format using msConvert (Chambers *et al.*, 2012). We processed these files using XCMS and AutoTuner (Smith *et al.*, 2006; Tautenhahn *et al.*, 2008; McLean & Kujawinski, 2020) to generate a spreadsheet of features. We define features as chromatographic peaks with unique mass-to-charge (*m/z*) and retention time values, with relative abundances determined by their area. We subjected processed data to quality-control filtering by removing possible contaminants and non-reproducible features as described previously (Dunn *et al.*, 2011). Briefly, we removed features within blanks, features with a coefficient of variation higher than 0.2 within pooled samples, and features with low reproducibility across factor groups. We report feature intensities normalized by the cell counts from day 3.

We used MAVEN to integrate compound peak areas within targeted data (Melamud *et al.*, 2010). We used an in-house MATLAB script to apply quality-control filtering and to quantify peak areas using a standard curve of 4 to 10 points. We retained metabolites for this analysis if the peak included a confirm ion, and the metabolite was present within two of three biological replicates for each treatment. We further culled the list by correcting for metabolite presence in procedural blanks.

### Statistics and Data Analysis

We used ANalysis Of VAriance (ANOVA) hypothesis testing to identify significantly different untargeted mass spectral features, targeted compound abundances, and growth time points. We identified significant pairwise-comparisons using Tukey’s honestly significant difference test (Tukey-HSD Test). We used linear models to identify significant trends between Hessa et al. (2005) and Wimley and White (1996) amino acid hydrophobicity scales and feature retention time. We applied Benjamini-Hochberg correction to control for type 1 error following all tests, and considered any *p-value* equal to or less than 0.05 to be significant.

We putatively annotated features to Kyoto Encyclopedia of Genes and Genomes (KEGG) compounds and tetrapeptides if feature masses were within 2.5 ppm error of the expected ion masses (Kanehisa, 2000). Tetrapeptide masses represent the set of all possible masses of any four amino acids linked together by a peptide bond. We used mummichog to match features to KEGG compounds (Li *et al.*, 2013). Whenever possible, we matched MS/MS spectra with *in silico* modeled MS/MS spectra of known compounds from MetFrag to increase strength of annotation (Ruttkies *et al.*, 2016). Due to prohibitive costs required to confirm all features with authentic standards, we focused on features pertinent to pathways involved in central metabolism and intracellular scavenging.

We obtained previously-published transcriptome data (see Table S1 from reference 15) from *H. akashiwo* strain CCMP 2393 grown under identical conditions to our study. We combined the expression of all reads mapping to a single KEGG ortholog to find the net expression of the putatively identified KEGG orthologs prior to analysis of the data. We used Analysis of Sequence Counts (ASC) to identify transcripts with a posterior *p-value* (post-*p*) > 0.95 for a log_2_ fold change greater than 2 or less than −2. All transcriptomic comparisons were between data from stress and replete cultures (Wu *et al.*, 2010). We considered individual transcripts satisfying either of these criteria to be significantly more or less abundant. When analyzing groups of transcripts together across nutrient treatments, we first normalized individual genes by the mean expression of that gene to remove baseline differences across genes. We applied the Wilcoxon-Test to check if a group of transcripts was significantly more or less abundant under a given stress condition relative to the control. We corrected all *p-values* for multiple comparisons using Bonferroni method. We considered any *p-value* equal to or less than 0.05 to be significant.

## Results and Discussion

Raphidophytes are ubiquitous in estuarine and coastal systems worldwide and their blooms cause severe damage to fisheries and local ecosystems. Yet, they are understudied relative to other marine phytoplankton groups like diatoms and coccolithophores. Here we used a combined metabolomic and transcriptomic approach to build a conceptual model of how the raphidophyte *H. akashiwo* remodels its metabolism under N- and P-stress, and compare that to other phytoplankton taxa.

### P-stressed cells catabolize lipids for sugar synthesis

Central carbon metabolism is the biochemical hub connecting intracellular macromolecular pools. Its net flux drives differences in intracellular nutrient stoichiometry (Voet & Voet, 2010). To understand its direction under P-stress, we first sought to determine whether glycolysis or gluconeogenesis was taking place. These pathways interconvert sugars and organic acids, and are distinguished by a few non-reversible reactions. Hence, we evaluated the differential expression of transcripts unique to each pathway (Fig 1a). Within P-stressed cells, two out of three gluconeogenic transcripts were significantly more abundant (post-*p* > 0.95, ASC, log_2_(FC) > 2), while one of the three glycolytic transcripts was significantly more abundant (post-*p* > 0.95, ASC, log_2_(FC) > 2), suggesting that P-stressed cells transcriptionally upregulate gluconeogenesis over glycolysis. The log_2_ fold change between the mean expression of gluconeogenic and glycolytic exclusive transcripts was 2.4, consistent with this hypothesis (Fig S2). To determine if increased gluconeogenic transcription resulted in a concomitant increase of pathway activity, we quantified trehalose, a gluconeogenic end member molecule. Within P-stressed cells, trehalose concentrations were significantly (*p* < 0.05, Tukey-HSD Test, n = 3) enriched (Fig 1b), supporting the transcript results. Based on these findings, we conclude that P-stress induces gluconeogenesis in *H. akashiwo*.

**Figure 1:**
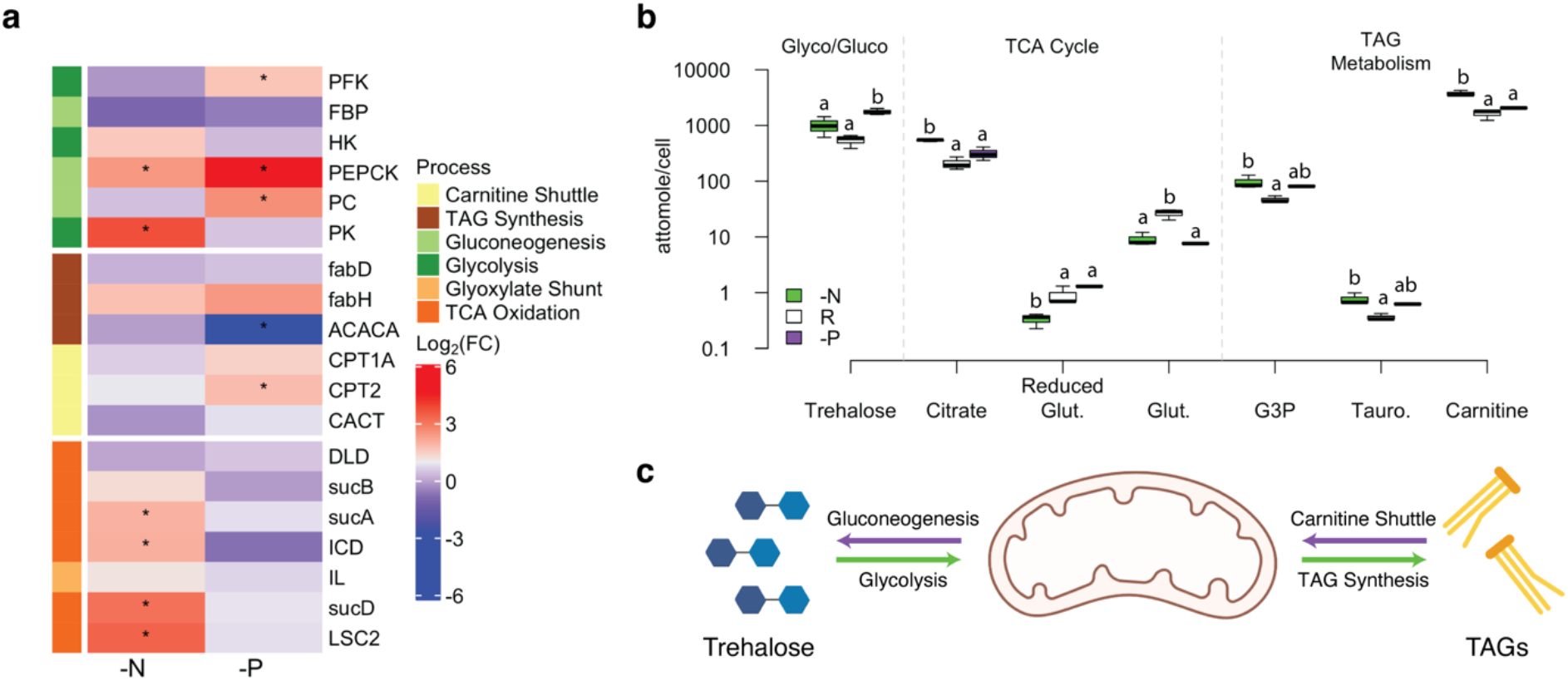
Central carbon metabolism stress response. A) Transcriptomic data on irreversible steps of gluconeogenesis, glycolysis, TCA oxidation, glyoxylate shunt, carnitine shuttle (CS), and triacylglyceride (TAG) lipid synthesis. Data are from N-stress or P-stress normalized to the condition in the replete cells (-N/R and -P/R, respectively). * - denotes significance defined as a log_2_(FC) > 2 and ASC post-p > 0.95. B) Metabolomics data supporting hypothesized pathway activity. We present data of individual metabolites in the order of N-stressed (-N, green), nutrient replete (R, white), and P-stressed (-P, purple) treatments. Boxes with distinct letters above them are significantly different as defined by pairwise Tukey-HSD test p < 0.05. C) Hypothesized fluxes of central carbon metabolism under N-stress (green) and P-stress (purple). Transcript abbreviations: PFK – phosphofructokinase, PK – pyruvate kinase, HK – hexose kinase, PEPCK – phosphoenolpyruvate carboxykinase, PC – pyruvate carboxylase, FBP – fructo-bisphosphate phosphatase, fabD – malonyl carrier protein transacylase, fabH – oxoacyl carrier protein synthase III, fabH – oxoacyl carrier protein synthase III, ACACA – acyl-CoA carboxylase, CPT1A – carnitine O-palmitoyltransferase 1, CPT2 – carnitine O-palmitoyltransferase 2, CACT – mitochondrial carnitine/acylcarnitine transporter, DLD – dihydrolipoamide dehydrogenase, sucB – 2-oxoglutarate dehydrogenase complex (dihydrolipoamide dehydrogenase), sucA – 2-oxoglutarate dehydrogenase complex (E1 component), ICD – isocitrate dehydrogenase, LSC2 – succinyl-CoA synthetase (beta subunit), sucD – succinyl-CoA synthetase (alpha subunit), IL – isocitrate lyase, FC – fold change. Metabolomic abbreviations: Glut. – glutathione, G3P – glycerol-3-phosphate, Tauro. – taurocholate.

Gluconeogenesis requires a flux of reduced carbon from the mitochondria. For reduced carbon to leave the mitochondria, it must avoid the tricarboxylic acid (TCA) cycle oxidation via the glyoxylate shunt (GS) (Ahn *et al.*, 2016). To determine whether GS or oxidation was more prevalent within P-stressed cells, we evaluated the differential expression of transcripts driving the bifurcation between the pathways; isocitrate lyase (IL) for GS and isocitrate dehydrogenase (ICD) for oxidation. IL had a log_2_ fold change of 0.74 while ICD had a fold change of −0.73, suggesting that P-stress transcriptionally upregulates GS over oxidation (Fig 1a). Cells oxidizing less carbon would experience less oxidative stress. To evaluate if this was the case for P-stressed cells, we calculated the ratio of reduced glutathione to glutathione, a measure of cellular oxidative stress (Dringen *et al.*, 2000). This ratio was significantly (*p* < 0.05, Tukey-HSD, n = 3) decreased in P-stressed cells, supporting our hypothesis (Fig S3b). Our combined metabolite and transcriptomic data imply that under P-stress, *H. akashiwo* engages the GS over carbon-oxidizing reactions.

Mitochondrial carbon bound for gluconeogenesis may originate from the catabolism of triacylglyceride (TAG) lipids (Voet & Voet, 2010). To be catabolized, TAGs must be removed from storage droplets with the assistance of cholic acid derivatives (Voet & Voet, 2010). Hence, we quantified taurocholate, a cholic acid derivative. While not significant (*p* < 0.07, Tukey-HSD Test, n = 3), P-stressed cells had elevated taurocholate concentrations suggesting that P-stress may enhance TAG mobilization (Fig 1b). We next sought to determine if the hypothesized increase in TAG mobility was due to production or catabolism of TAGs. For this, we evaluated the differential expression of transcripts from both the carnitine shuttle (CS) and cytosolic lipid elongation pathways. The carnitine shuttle is considered the rate limiting step of lipid catabolism (Houten & Wanders, 2010) and is regulated to oppose lipid elongation to avoid futile cycles (Voet & Voet, 2010). We observed that one out of three carnitine shuttle transcripts were significantly more abundant (post-*p* > 0.95, ASC, log_2_(FC) > 2) within P-stressed cells. By contrast, one out of three lipid elongation enzymes was significantly less abundant (post-*p* > 0.95, ASC, log_2_(FC) > 2) within P-stressed cells. In order to reinforce these findings, we calculated a log_2_ fold change between the averaged expression of carnitine shuttle and lipid elongation exclusive transcripts, which showed a value of 0.18, corroborating our hypothesis (Fig S2). These dynamics suggest that TAG catabolism is being upregulated by P-stressed cells (Fig 1a). We sought to confirm evidence of carnitine shuttle activity by quantifying carnitine but its concentration was not significantly different in P-stressed cells than replete ones (Fig 1b). One possible explanation is that in preparation for TAG degradation, P-stressed cells accumulate acyl-carnitines (Kong *et al.*, 2018). These molecules were not detected by our analytical method and future small molecule quantification experiments are needed to confirm this hypothesis. Additional targeted lipidomic analysis measuring TAG concentrations would serve to validate our hypothesized trends.

These concerted pathway level changes suggest that under P-stress, *H. akashiwo* drives gluconeogenesis via the catabolism of TAG carbon (Fig 1c). The system-level dynamics are most similar to those of P-stressed diatoms, as these organisms are hypothesized to upregulate gluconeogenesis and the carnitine shuttle under P-stress (Brembu *et al.*, 2017; Alipanah *et al.*, 2018). By contrast, P-stressed metabolism in the coccolithophore, *Emiliania huxleyi,* appears to be quite different. Prior studies reported that this organism upregulates gluconeogenesis, glycolysis, and TAG synthesis in tandem (Rokitta *et al.*, 2016; Wördenweber *et al.*, 2018). To our knowledge, this is the first report of P-stress driven glyoxylate shuttle transcription and trehalose enrichment among phytoplankton groups. These distinguishing features may underlie physiological differences exhibited between *H. akashiwo* and other closely related phytoplankton.

### N-stressed cells increase respiration and store excess carbon as lipids

Like our analysis of P-stressed cells, we sought to determine the direction of central carbon metabolism in N-stressed cells. Again, we evaluated whether glycolysis or gluconeogenesis was upregulated by examining the differential expression of pathway-specific transcripts. The results were equivocal, with one of three gluconeogenic transcripts significantly more abundant (post-*p* > 0.95, ASC, log_2_(FC) > 2) within N-stressed cells, and one of three glycolytic transcripts significantly more abundant (post-*p* > 0.95, ASC, log_2_(FC) > 2) within N-stressed cells, suggesting either pathway may be taking place (Fig 1a). It is possible that our calculation integrates gene expression from versions of the pathway that are localized to the cytosol and the plastid, which may be differentially regulated under N-stress (Brembu *et al.*, 2017). Constraining the cellular location is not possible due to the lack of a fully sequenced genome for *H. akashiwo*. In order to differentiate the activity between pathways, we calculated a log_2_ fold change of −1.2 between the averaged expression of gluconeogenic relative to glycolytic exclusive transcripts, suggesting that gluconeogenic transcripts are less abundant than glycolytic ones within N-stressed cells (Fig S2). In addition, trehalose concentrations were not significantly enriched (Fig 1b), pointing to lower gluconeogenic activity in N-stressed cells. These combined findings suggest that N-stressed cells favor glycolysis over gluconeogenesis.

Increased glycolytic carbon flux would foster greater TCA cycle oxidation. To determine if oxidation was preferentially upregulated in N-stressed cells, we examined the differential expression and the log_2_ fold change of IL to ICD. We observed that ICD was significantly more abundant (post-*p* > 0.95, ASC, log_2_(FC) > 2) under N-stress (Fig 1a) and the ratio of IL to ICD had a log_2_ fold change of −0.98, supporting the hypothesis that N-stressed cells increase carbon oxidation (Fig S2). We then evaluated the differential expression of transcripts downstream of ICD driving oxidation reactions. We observed that three out of five transcripts were significantly more abundant (post-*p* > 0.95, ASC, log_2_(FC) > 2), supporting our hypothesis of increased oxidation (Fig 1a). To evaluate if transcriptional upregulation patterns resulted in a tandem increase in TCA cycle activity, we quantified citrate; citrate accumulation is a common signature of upregulated TCA cycle flux (Voet & Voet, 2010). We observed that citrate concentrations were significantly higher (*p* < 0.05, Tukey-HSD Test, n = 3) under N-stress, consistent with our hypothesis of increased TCA cycling (Fig 1b). To determine if N-stressed cells experienced elevated oxidative stress from TCA cycling, we evaluated glutathione recycling. Although we observed that the ratio of reduced glutathione to glutathione was not significantly different under N-stress, the concentration of reduced glutathione was significantly depleted (*p* < 0.05, Tukey-HSD Test, n = 3) (Fig 1b) and transcripts driving glutathione recycling were significantly more abundant (*p* < 0.05, Wilcoxon-test, n = 6) (Fig S3a). Reduced glutathione is an N-rich molecule; hence its observed depletion may be due to N reallocation. The increased transcription of glutathione recycling enzymes may suggest that N-stressed cells overcome the reduction in reduced glutathione concentrations by increasing its recycling rates. These findings support the hypothesis that N-stressed cells upregulate TCA cycle carbon oxidation.

Increased TCA cycling oxidation often results in TAG enrichment. Prior studies show that N-stressed *H. akashiwo* cells are enriched with TAG lipids (Stewart *et al.*, 2015; Bianco *et al.*, 2016). In our dataset, concentrations of taurocholate, a TAG mobilization marker, were significantly higher (*p* < 0.05, Tukey-HSD Test, n = 3) within N-stressed cells, suggesting elevated TAG mobilization under N-stress (Fig 1b). To determine if mobilization supporting the storage of newly synthesized TAGs, we quantified glycerol-3-phosphate, a TAG lipid backbone, which was significantly higher (*p* < 0.05, Tukey-HSD Test, n = 3) in N-stressed cells (Fig 1b). We next investigated changes in transcription of TAG synthesis genes. Specifically, we evaluated the transcriptome changes of the carnitine shuttle and enzymes initiating lipid elongation. While none of these transcripts were significantly more or less abundant, the log_2_ fold change of the averaged expression of carnitine shuttle and lipid elongation exclusive transcripts showed a value of −0.38, suggesting that elongation was slightly upregulated relative to the carnitine shuttle (Fig S2). Although we did not measure TAGs within this study, our results suggest elevated TAG production due to enrichment of precursors and depletion of TAG catabolism transcripts.

These concerted pathway-level changes suggest that N-stress drives TAG synthesis via an increase in glycolysis (Fig 1c). Our findings match previously-published isotope tracer experiments on N-stressed *H. akashiwo* (Takahashi & Ikagawa, 1988). Similar observations of increased glycolysis, TCA cycling, and TAG synthesis have been reported for both N-stressed diatoms and *E. huxleyi* (Bender *et al.*, 2014; Rokitta *et al.*, 2014; Alipanah *et al.*, 2015; Kim *et al.*, 2017). Unlike diatoms and *E. huxleyi, H. akashiwo* enriches citrate (Bromke *et al.*, 2015; Wördenweber *et al.*, 2018), which can allosterically inhibit glycolysis or increase TAG synthesis rates (Voet & Voet, 2010). Citrate enrichment may indicate that *H. akashiwo* differs from other phytoplankton in these two processes.

N-stress has been reported to be linked to the initiation of the diel migration employed by *H. akashiwo* during blooms (Ji *et al.*, 2018; Ji *et al.*, 2020). The diel migration is hypothesized to serve *H. akashiwo* by helping it acquire dissolved nutrients below the thermocline (Smayda, 1998). Sinking rates are linked to TAG accumulation and the resulting increase in cellular specific gravity (Hatano *et al.*, 1983; Wada *et al.*, 1985; Wada *et al.*, 1987). Hence, TAG synthesis *in situ* may be driven by an increased in glycolysis and TCA cycle oxidation as presented here for N-stressed cells. The system level dynamics described here should be considered when building models of *H. akashiwo* blooms.

### Nutrient stress responses use central metabolism in opposite ways

*H. akashiwo* appears to use central carbon metabolism in two distinct ways to overcome N- and P-stress, in contrast to *E. huxleyi*, which responds to N- and P-stress similarly (Rokitta *et al.*, 2016; Wördenweber *et al.*, 2018). Our hypothesis of stress-specific metabolic dynamics is based on the enrichment and proposed sources of pathway end-member storage molecules, trehalose and TAGs. These end-member molecules vary drastically in their potential to contribute towards future biomass and other cellular functions. Trehalose can enter glycolysis after one reaction (breakage of the disaccharide bond), and may be quickly redirected towards the synthesis of nucleic and amino acids (Voet & Voet, 2010). In contrast, TAGs must be converted into sugars via gluconeogenesis before this is possible. However, TAGs provide far higher amounts of ATP per carbon than trehalose. It is critical to consider these trade-offs when building models to describe *H. akashiwo* ecosystem processes, as accumulation of either would result in distinct impacts on fitness (Sharma & Steuer, 2019). As TAGs are far more carbon-rich than trehalose, the differences in orientation of central carbon metabolism may also explain why N-stressed cells had a greater measured C:N (14.37) ratio than that (8.95) of P-stressed cells (Haley *et al.*, 2017). These metabolic nuances and their hypothesized physiological consequences speak to the importance of understanding the metabolism of more non-model phytoplankton to develop models to characterize bloom and nutrient cycling dynamics (Worden *et al.*, 2015). These findings would not have been possible without the tandem analysis of transcriptomic and metabolomic data (Wördenweber *et al.*, 2018). Hence, our approach may serve investigators of raphidophytes and other non-model phytoplankton in similar ways.

### Intracellular recycling is pervasive under N-stress

Phytoplankton, including *H. akashiwo*, employ various strategies in response to nutrient stress (Van Mooy *et al.*, 2009; Allen *et al.*, 2011; Grzymski & Dussaq, 2012; Harke *et al.*, 2017; Kujawinski *et al.*, 2017; Read *et al.*, 2017). A prior transcriptomic-based study showed that under N- and P-stress, *H. akashiwo* increases extracellular inorganic nutrient scavenging transporters, recycles amino acid N via the urea cycle, and catabolizes photosynthetic machinery (Haley *et al.*, 2017). However, transcriptome studies alone are unable to confirm these processes in most regards without metabolite data as an end member to the biological cascade initiated by transcription. For example, prior work with *H. akashiwo* was unable to identify which amino acids drive the urea cycle or if additional intracellular recycling processes enabled the observed upregulation of over 30% of the transcriptome under N-stress (Haley *et al.*, 2017). Here, we evaluated our data to uncover possible intracellular macromolecular recycling processes.

Previously, it was shown that urea cycle transcripts are significantly more abundant within N-stressed *H. akashiwo* cells (Haley *et al.*, 2017). Indeed, our data also show that urea cycle intermediates are enriched within N-stressed cells, confirming these trends (see Note S2 and Fig S4). However, the source of these compounds is unclear and could include either extracellular uptake or the degradation of endogenous protein. Prior studies show that *H. akashiwo* favors uptake of inorganic N over organic N relative to sympatric phytoplankton (Zhang *et al.*, 2019). Hence, we hypothesized that *H. akashiwo* may sustain increased urea cycling by degrading endogenous proteins.

One endogenous protein degradation mechanism is proteasome-mediated enzyme degradation (PMED) (Boer *et al.*, 2010). PMED is an ATP-dependent process where enzymes are tagged with ubiquitin, shuttled into the proteasome, and released as peptides between 4 and 20 amino acids in length (Fig 2a). We observed that putatively-identified tetrapeptides from untargeted data were significantly depleted (*p* < 10^−10^, Kruskal-Wallis Test, n = 597) under both P- and N-stress, suggesting differential activity of PMED in both stress cultures (see Note S3). To evaluate whether differential PMED activity was up- or down-regulated under stress, we gathered transcripts involved in enzyme ubiquitin tagging (ubiquination), proteasome biosynthesis, and downstream peptide cleavage (peptidases). We observed that transcripts for eight out of ten sub processes within PMED were significantly more abundant (*p* < 0.05, Wilcoxon-Test) in N-stressed cells, while three out of ten were significantly more abundant (*p* < 0.05, Wilcoxon-Test) and one was significantly less abundant (p < 0.05, Wilcoxon-Test) in P-stressed cells (Fig 2b). To test whether this transcriptional upregulation corresponded with an increase in peptide degradation activity, we quantified hydroxyproline. Hydroxyproline is synthesized via post-translational modifications of proteins (Allen *et al.*, 2008), hence its cytosolic concentration serves as evidence of protein degradation. Hydroxyproline concentrations were significantly enriched (*p* < 0.05, Tukey-HSD Test, n = 3) in N-stressed cells exclusively. Our paired transcriptomic and metabolomics datasets suggest that *H. akashiwo* upregulates proteasome enzyme degradation to overcome N-stress. To our knowledge, this is the first indicator of PMED as an N-stress mitigation strategy in phytoplankton. PMED is highly specific, hence it may contribute to an observed proteome-level depletion of N-rich proteins, as observed in the N-stressed green alga *Chlamydomonas reinhardtii* (Schmollinger *et al.*, 2014). In addition to amino acid recycling, we sought to constrain if nucleotide recycling enabled a reallocation of N within N-stressed *H. akashiwo*. To check this hypothesis, we first quantified 17 distinct intermediates within nucleic acid metabolism (Fig 3a). Surprisingly, we observed an enrichment of nucleic acid bases and nucleosides and a significant depletion of nucleotide monophosphates (NMPs) (Fig 3a). Indeed, five out of eleven measured nucleoside or nucleic acid bases were significantly enriched (*p* < 0.05, Tukey-HSD, n = 3) under N-stress. Similar metabolite trends were reported in N-stressed yeast due to the autophagy-mediated breakdown of ribosomes and other nucleic acids (Xu *et al.*, 2013). To investigate if this mechanism could support metabolite enrichments, we gathered the gene expression of 21 DNA and/or RNA degradation enzymes (Fig 3b). These transcripts were significantly more abundant within N-stressed cells (*p* < 10^−10^, Wilcoxon Test, n = 21), suggesting that autophagic breakdown may drive our observed metabolite trends. Unfortunately, none of our annotated transcripts corresponded to reactions driving interconversion reactions between distinct purines or pyrimidines. This may be due to the annotation challenges, as studies of this pathway in yeast faced similar obstacles (Xu *et al.*, 2013).

**Figure 2:**
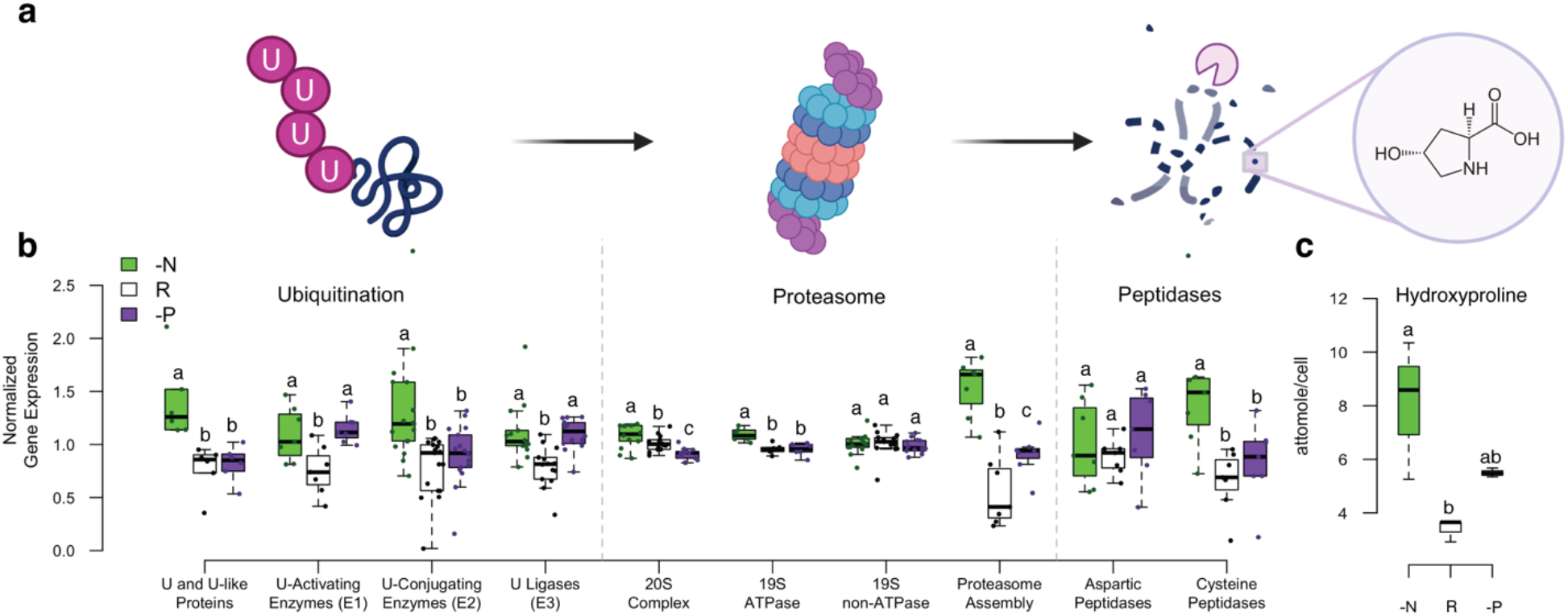
Proteasome mediated enzyme degradation. A) Schematic overview of proteasome enzyme degradation process. Enzymes tagged with ubiquitin (U) are brought into the proteasome to be broken down into peptides between 4 and 20 amino acids in length. Peptides are then broken down by cytosolic peptidases. B) Normalized gene expression values for distinct processes related to enzyme ubiquitination, the proteasome, and cytosolic peptidases. Genes were normalized by the mean expression of enzymes across all treatments. Boxes with distinct letters above them are significantly different as defined by pairwise Wilcoxon Test and Bonferroni correction p < 0.05. C) Hydroxyproline concentrations, an enzyme degradation by-product. Boxes with distinct letters above them are significantly different as defined by pairwise Tukey-HSD test p < 0.05. Abbreviations: -N - N-stressed, R - nutrient replete, -P - P-stressed, U – ubiquitin.

**Figure 3:**
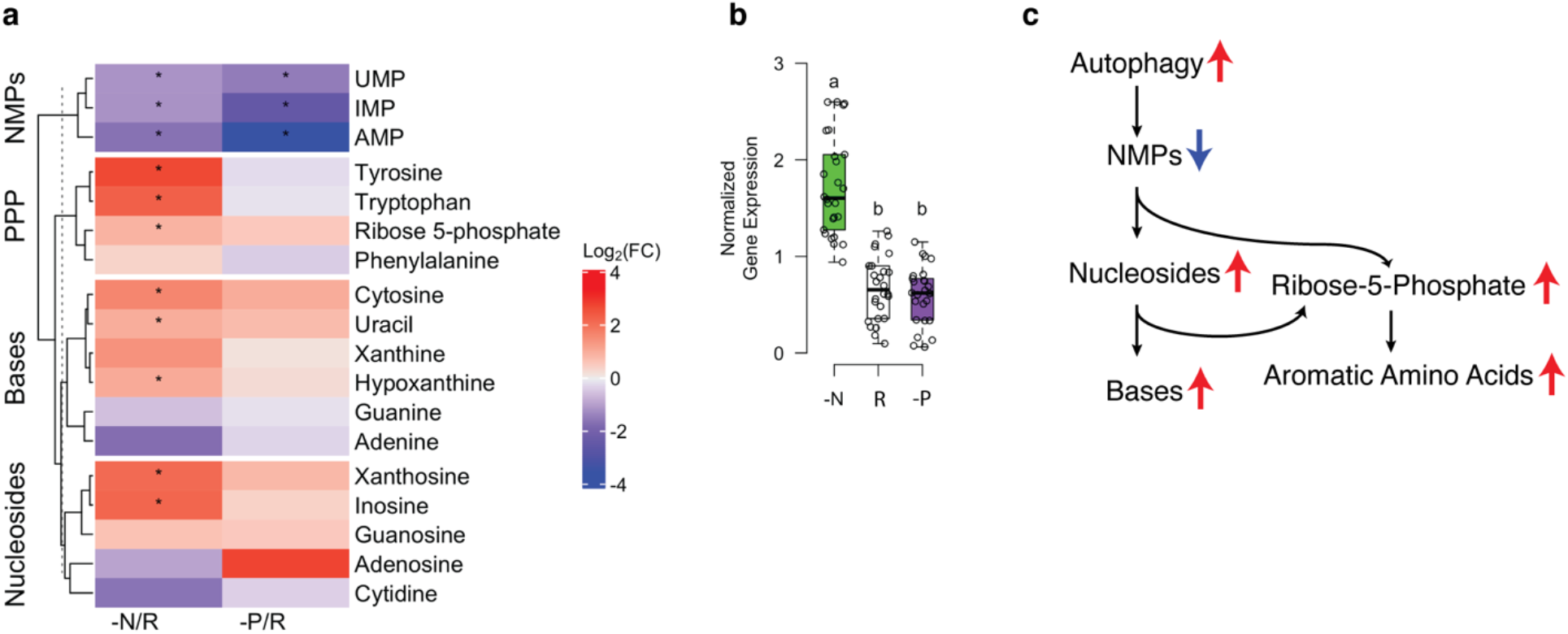
Stress induced nucleic acid scavenging. A) Targeted metabolite data of nucleoside monophosphates (NMPs), nucleic acid bases (Bases), nucleosides, and pentose phosphate pathway and aromatic amino acids (PPP). * denotes significant enrichment or depletion of compound concentration (p < 0.05, Tukey-HSD Test, n = 3). B) Normalized gene expression for RNA and DNA cleavage enzymes. Boxes with distinct letters above them are significantly different as defined by pairwise Wilcoxon Test and Bonferroni correction p < 0.05. C) Proposed pathway dynamics within N-stressed cells. Colored arrows adjacent to names indicate the net enrichment/depletion status of compounds or processes described by our data. Abbreviations: -N - N-stressed, R - nutrient replete, -P - P-stressed, UMP – uridine monophosphate, IMP – inosine monophosphate, AMP – adenosine monophosphate.

To determine if nucleic acid recycling within *H. akashiwo* produced a downstream response similar to yeast, we evaluated the activity of the pentose phosphate pathway (PPP). In yeast, carbon scavenged from NMPs leads to both an enrichment of PPP metabolite ribose-5-phosphate and an increase in non-oxidative PPP activity (Xu *et al.*, 2013). We quantified ribose-5-phophate, and observed that ribose-5-phosphate concentrations were significantly higher (*p* < 0.05, Tukey-HSD, n = 3) under N-stress (Fig 3a), similar to yeast. However, when we gathered transcripts for non-oxidative PPP reactions, we observed that three out of four non-oxidative PPP transcripts were significantly less abundant (post-*p* > 0.95, ASC, log_2_(FC) > 2) within N-stressed cells, contrary to our expectation (Fig S5). This result suggests that ribose-5-phosphate derived from nucleic acids degradation may support an alternative function other than non-oxidative PPP activity within *H. akashiwo*.

One possible alternative is the biosynthesis of aromatic amino acids. These compounds all originate from erythrose-4-phosphate, a downstream product of ribose-5-phosphate (Voet & Voet, 2010). We evaluated the activity of the aromatic amino acid biosynthesis pathway (Fig 3a and Fig S5) and observed that two out of ten transcripts were significantly more abundant (post-*p* > 0.95, ASC, log_2_(FC) > 2) and that transcription of enzymes within the pathway was enriched, although not statistically significant (*p* < 0.06, Wilcoxon-Test, n = 12), in N-stressed cells. Additionally, all aromatic amino acid biosynthesis metabolites with the exception of phenylalanine were significantly enriched (*p* < 0.05, Tukey-HSD Test, n = 3) in N-stressed cells (Fig 3a). Taken together, these observations suggest that ribose scavenged from NMP degradation drives aromatic amino acid biosynthesis (Fig 3c). Within plants and algae, most aromatic amino acids serve as building blocks for specialized metabolites, from electron carriers to natural products and chemical signals (Lynch & Dudareva, 2020). For example, tryptophan secreted by the diatom *Thalassiosira pseudonana* was shown to drive mutualistic growth between itself and a sympatric microbe (Amin *et al.*, 2015). *H. akashiwo* may rely on a similar strategy to overcome N-stress. Studies on diatoms have noted that nutrient stress increases PPP activity (Alipanah *et al.*, 2018). However, to our knowledge, we provide the first evidence of either the connection between the PPP and nucleic acid degradation or its connection to aromatic amino acid biosynthesis. Nucleoside enrichment patterns have been observed in other marine phytoplankton, as inosine was also enriched in N-stressed *Prochlorococcus* (Szul *et al.*, 2019). Hence, similar metabolic dynamics may support other phytoplankton.

Intracellular recycling appears to be a critical nutrient N-stress response strategy within *H. akashiwo*. Recycling may support *H. akashiwo* life history and behavior changes such as diel migration in addition to supporting sustained growth in resource limiting situations. Modeling of *H. akashiwo* will need to consider intracellular recycling and its impact on fitness for predicting behavior and bloom dynamics. The extent to which these processes uniquely define the niche of *H. akashiwo* or other raphidophytes, relative to other phytoplankton is uncertain and underscores the need for additional multi-‘omic studies across a range of phytoplankton lineages.

### Phytoplankton nutrient stress biomarkers reveal stress status within *H. akashiwo*

Our data allowed us to evaluate the efficacy of proposed phytoplankton N-stress (glutamine:glutamate) and P-stress (adenosine monophosphate:adenosine) diagnostics within *H. akashiwo* (Boer *et al.*, 2010; Kujawinski *et al.*, 2017). We observed that N-stressed cells had significantly lower (*p* < 0.05, Tukey-HSD Test, n = 3) glutamine-to-glutamate ratios relative to the other treatments, while the P-stressed cells had significantly lower (*p* < 0.05, Tukey-HSD Test, n = 3) AMP-to-adenosine ratios (Fig 4). To our knowledge, this is the first evidence of the utility of these stress diagnostics in any raphidophyte. Measurements of these ratios in field raphidophyte populations would serve as an important new approach for identifying whether a population is experiencing N- or P-stress, and defining the resource controls on bloom dynamics *in situ*.

**Figure 4:**
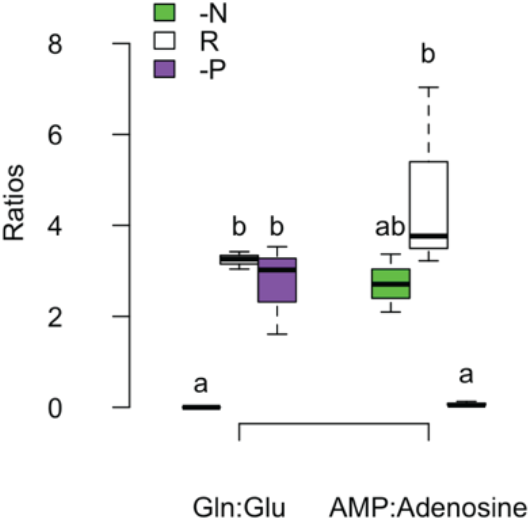
Diagnostic N-stress and P-stress ratios in H. akashiwo. Ratios were calculated using measured concentrations of each metabolite. Boxes with distinct letters above them are significantly different as defined by pairwise Tukey-HSD test p < 0.05. Abbreviations: Gln – glutamine, Glu – glutamic acid, AMP – adenosine monophosphate.

## Conclusions

This work employs a multi-omics approach to explore the impact of N- and P-stress on the HAB-forming raphidophyte, *H. akashiwo*. We characterized the stress-mediated system-level changes within central carbon metabolism and revealed that intracellular recycling of macromolecules is pervasive under stress (Fig 5). Under N- and P-stress, *H. akashiwo* showed similar central carbon metabolism acclimation patterns as other model phytoplankton under nutrient stress. However, fine scale enrichment of distinct molecules distinguished its metabolic shifts from more frequently-studied phytoplankton (Fig 5). Identifying these differences would not have been possible without a multi-‘omics approach. Similarly, evidence of novel intracellular recycling pathways could support sustained growth of *H. akashiwo* under conditions of low N and may be a mechanism underpinning niche segregation among competitors more reliant on N uptake. Taken together, these insights suggest that nutrient stress has distinct physiological impacts on *H. akashiwo* relative to other model phytoplankton, like diatoms. Our results suggest metabolism data from model phytoplankton does not capture the nuances of non-model phytoplankton completely. Hence, more characterizations of metabolic stress responses within other non-model phytoplankton are critical to accurately predict, manage, and prevent HABs, and understand phytoplankton community composition and function in a changing ocean (Worden *et al.*, 2015).

**Figure 5:**
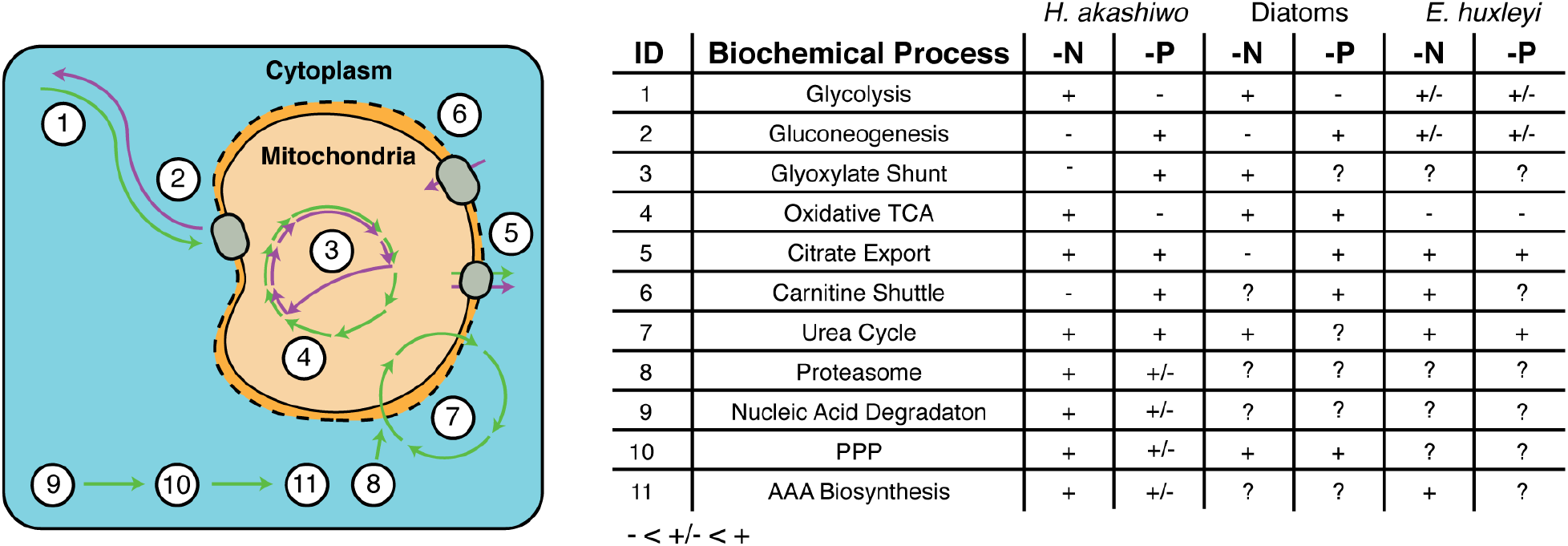
Conceptual model summarizing the changes in metabolism of H. akashiwo under nutrient stress. Each white numbered circle in the model displays a distinct intracellular process responding to stress. The purple and green arrows denote predicted systems-level biochemical activity for P-stressed and N-stressed H. akashiwo, respectively. + and − signs indicate increased and decreased activity, respectively, in each phenotype relative to the replete phenotype. Values for diatoms and E. huxleyi are based on reported pathway enrichment trends from prior studies (Rokitta et al., 2014; Alipanah et al., 2015; Rokitta et al., 2016; Brembu et al., 2017; Alipanah et al., 2018; Wördenweber et al., 2018). Boxes with question marks denote biochemical processes without existing data. Diatoms includes data from T. pseudonana, Phaeodactylum tricornutum, and Skeletonema costatum. Abbreviations: PPP – pentose phosphate pathway, AAA – aromatic amino acid.

## Acknowledgements

We thank Krista Longnecker for her helpful contributions and constructive review of the manuscript, and Penny Chisholm, Martin Polz, and Ben Temperton for feedback on analysis and results. Funding support included the National GEM Consortium and the NSF graduate research fellowship program fellowships (CM) and grants from the Gordon and Betty Moore Foundation (#3304 to EBK) and the Simons Foundation (#509034 to EBK, SCOPE award ID 329108 and ID 721225 to STD). Support was also provided by the Paul M. Angell Family Foundation and the Vetlesen Foundation to STD. The transcriptome samples analyzed in this study are MMETSP0292, MMETSP0294, and MMETSP0295 which were sequenced, assembled, and annotated with the ABySS pipeline at the National Center for Genome Resources with support from the Gordon and Betty Moore Foundation through Grant #2637 to the National Center for Genome Resources. These samples were submitted by Andrew Juhl and STD to the Marine Microbial Eukaryote Transcriptome Sequencing Project. Metabolite samples were analyzed in the WHOI FT-MS Facility, funded by grants from the National Science Foundation (OCE-0619608 to EBK and OCE-1058448 to EBK and MCKS). Figures of biological schematics were created using BioRender.com.

## Author Contribution

CM, EBK and STD designed the research and wrote the manuscript. CM analyzed results. CM and GJS performed chemical extractions. MCKS optimized mass spectrometry methods. STH grew *H. akashiwo* cultures. All authors contributed to the writing of the manuscript.

## Data Availability

Metabolomics data will be available within metabolights accession upon repository approval.

## Abbreviations

GS: glyoxylate shunt
TCA: tricarboxylic acid cycle
N: nitrogen
P: phosphorus
HAB: harmful algal blooms
TAG: triacylglyceride lipids
IL: isocitrate lyase
ICD: isocitrate dehydrogenase
NMP: nucleotide monophosphate
PPP: pentose phosphate pathway

